# Fork Slowing and Reversal as an Adaptive Response to Chronic ATR Inhibition

**DOI:** 10.1101/2021.05.18.444697

**Authors:** Diego Dibitetto, Andrea Sanchi, Ethan J. Sanford, Massimo Lopes, Marcus B. Smolka

**Affiliations:** Department of Molecular Biology and Genetics, Weill Institute for Cell and Molecular Biology, Cornell University, Ithaca, NY 14853, USA; Institute of Molecular Cancer Research, University of Zurich, 8057 Zurich, Switzerland

## Abstract

Inhibitors of the replication stress response kinase ATR are currently being explored in anti-cancer therapy. Acute ATR inhibition is known to impair the proper control of origin firing, DNA repair, and cell cycle, resulting in DNA breaks and mitotic catastrophe. Less is understood about the effects of clinically relevant regimes of ATR inhibition, which involve chronic and low doses of ATR inhibitors (*cATRi*) to cells. Here we report unexpected molecular effects of *cATRi* on replication dynamics. *cATRi* strongly reduces fork speed but has minimal effects on the accumulation of DNA breaks or cell survival. *cATRi* promotes extensive fork reversal and RAD51- and PARP-mediated fork slowing that correlate with the accumulation of DNA-RNA hybrids. Our work shows that fork reversal is a critical adaptive response ensuring cell survival during *cATRi* and that the manipulation of fork reversal causes hypersensitivity to *cATRi*, increasing the effectiveness of ATR inhibitors in anti-cancer therapies.

## INTRODUCTION

Faithful replication of the genetic material is crucial for genome stability and cellular proliferation. Many cancers exhibit mis-regulation of DNA replication and become increasingly dependent on mechanisms for coping with chronic replication stress. A first responder to replicative stress is the ATR kinase (Saldivar et al., 2017), which senses a range of replication intermediates through the recognition of RPA bound to ssDNA (Byun et al., 2005; Shiotani and Zou, 2009; You et al., 2002; Zou and Elledge, 2003), and coordinates the firing of replication origins with the progression through the cell cycle (Buisson et al., 2015; Costanzo et al., 2003; Moiseeva et al., 2017, 2019; Saldivar et al., 2018; Toledo et al., 2013). ATR’s role in restraining origin firing mainly depends on the activation of CHK1, which limits CDK activity (Moiseeva et al., 2019; Petermann et al., 2010; Sanjiv et al., 2016). ATR also has key roles in rescuing collapsed forks by coordinating the action of homologous recombination (HR) factors (Lanz et al., 2019). Given the central role of ATR in protecting against oncogene-induced replication stress (Lecona and Fernandez-Capetillo, 2018), inhibition of ATR has been proposed to be an effective approach for selectively targeting cancer cells (Charrier et al., 2011; Toledo et al., 2011). Several preclinical studies showed that treatment with ATR inhibitors (ATRi) strongly synergizes with PARP inhibitors (PARPi), FDA-approved drugs used in the treatment of HR-deficient cancers (Barnieh et al., 2021; Dibitetto et al., 2020; Kim et al., 2018, 2017, 2020; Lloyd et al., 2020; Schoonen et al., 2019; Yazinski et al., 2017), and a range of clinical trials are currently underway. However, most of our mechanistic understanding of the impact of ATRi on cells derives from studies using acute treatment with high doses (micromolar range) of ATRi, which do not properly represent the therapeutic context used in clinical trials that rely on multiple doses of ATRi at sufficiently low concentrations to avoid serious contraindications for patients (www.clinicaltrials.gov). Importantly, our recent works revealed that a chronic ATRi treatment, which more closely mimics the therapeutic context of ATRi, results in outcomes not observed in response to acute ATRi treatments (Dibitetto et al., 2020; Kim et al., 2018). For example, while chronic ATRi results in a strong reduction in the abundance of HR factors, leading to reduced DNA end resection and HR capacity, acute ATRi did not have the same effect on HR factor abundance and did not impact resection (Dibitetto et al., 2020; Kim et al., 2018). Thus, to better understand how ATRi can be rationally and effectively used in clinical settings it is essential to thoroughly characterize the effects of chronic ATRi delivery.

In this study, we find that a clinically relevant regime for chronic delivery of the ATRi AZD6738 (referred here as *cATRi*), has a profound effect on fork dynamics in proliferating cells without causing the substantial genotoxicity that is observed upon acute ATRi. Analysis of fork architecture following *cATRi* shows that fork slowing is associated with a striking accumulation of reversed forks. Destabilization of fork reversal by PARPi upon *cATRi* results in increased fork speed, chromosomal breakage, and impaired cellular proliferation, shedding new lights into the mechanisms of the ATRi/PARPi synergy. Moreover, we found that DNA-RNA hybrids accumulate upon *cATRi* and that their forced resolution rescues fork speed. We propose that fork slowing and reversal are part of an adaptive cellular response that is crucial to maintain replisome stability and ensure cell survival following *cATRi*.

## RESULTS

### *cATRi* uncouples origin firing and fork speed from genomic instability

The current state of knowledge of how ATRi affects cancer cells comes largely from studies using acute delivery of micromolar concentrations of ATRi. To study the effects of an ATRi delivery regime that more closely recapitulates a therapy context, we used a chronic delivery of sub-micromolar concentrations of the ATRi AZD6738 (Foote et al., 2018), which is currently being used in several pre-clinical trials (Barnieh et al., 2021). Although the precise pharmacokinetic properties of AZD6738 are currently being assessed in patients, studies in mice have detected sub-micromolar concentrations in plasma samples after drug peritoneal injection or oral gavage (Checkley et al., 2015; Fròsina et al., 2018; Kiesel et al., 2017). These concentrations are minimally toxic to somatic cells but still sensitized cancer-derived xenografts to various chemotherapeutic agents (Kim et al., 2017; Vendetti et al., 2015; Wallez et al., 2018). We subjected U-2OS cells to a 5-day chronic treatment with a 0.4 µM AZD6738 concentration (hereafter called *cATRi*) and observed that *cATRi* did not adversely affect cellular proliferation after five days (Figure 1A). To better understand the impact of *cATRi* on DNA replication dynamics, we monitored fork progression using an established single-molecule DNA fiber assay after *in vivo* incorporation of thymidine analogs (Quinet et al., 2017). Analysis of replication tracts showed that *cATRi* drastically reduces fork elongation rate in U-2OS and HCT116 cells (Figure 1B). Consistent with previous reports showing a role for ATR in repressing origin firing (Buisson et al., 2015; Couch et al., 2013; Moiseeva et al., 2017, 2019; Mutreja et al., 2018; Rainey et al., 2020), we found that fork speed was rescued by CDC7 inhibition (Figure 1C). Increased origin firing upon acute ATRi is thought to generate DNA breaks and replication catastrophe in response to HU or aphidicolin (Couch et al., 2013; Toledo et al., 2013). Given the strong reduction in fork speed following *cATRi*, we next asked whether replication forks were more prone to breakage. We monitored DSB signaling markers upon *cATRi* and in cells after ionizing radiation (1 hour after 5 Gy) (Figure 1D). Interestingly, in contrast with irradiated cells, *cATRi* did not induce DSB signaling as analyzed by western blotting (Figure 1D). In particular, the phosphorylation pattern triggered by *cATRi* reflects a replication stress response involving phosphorylation of RPA2, and DNA-PKcs but not ATM (Figure 1D). Consistently, Neutral Comet assay results showed chromosome fragmentation in cells after ionizing radiation but not upon *cATRi* (Figure 1E). Moreover, we directly compared DSB signaling markers in cells following either *cATRi* or acute ATRi (*aATRi*), selecting a AZD6738 dose that was recently used to show a role of ATR in origin firing regulation (Moiseeva et al., 2019; Rainey et al., 2020). Consistent with our results showing the lack of extensive breaks upon *cATRi*, the signaling response induced by *cATRi* was markedly different from the response induced by *aATRi*, which confirmed the expected effect of *aATRi* in causing DSBs (Figure 1F) (Buisson et al., 2015).

**Figure 1.**
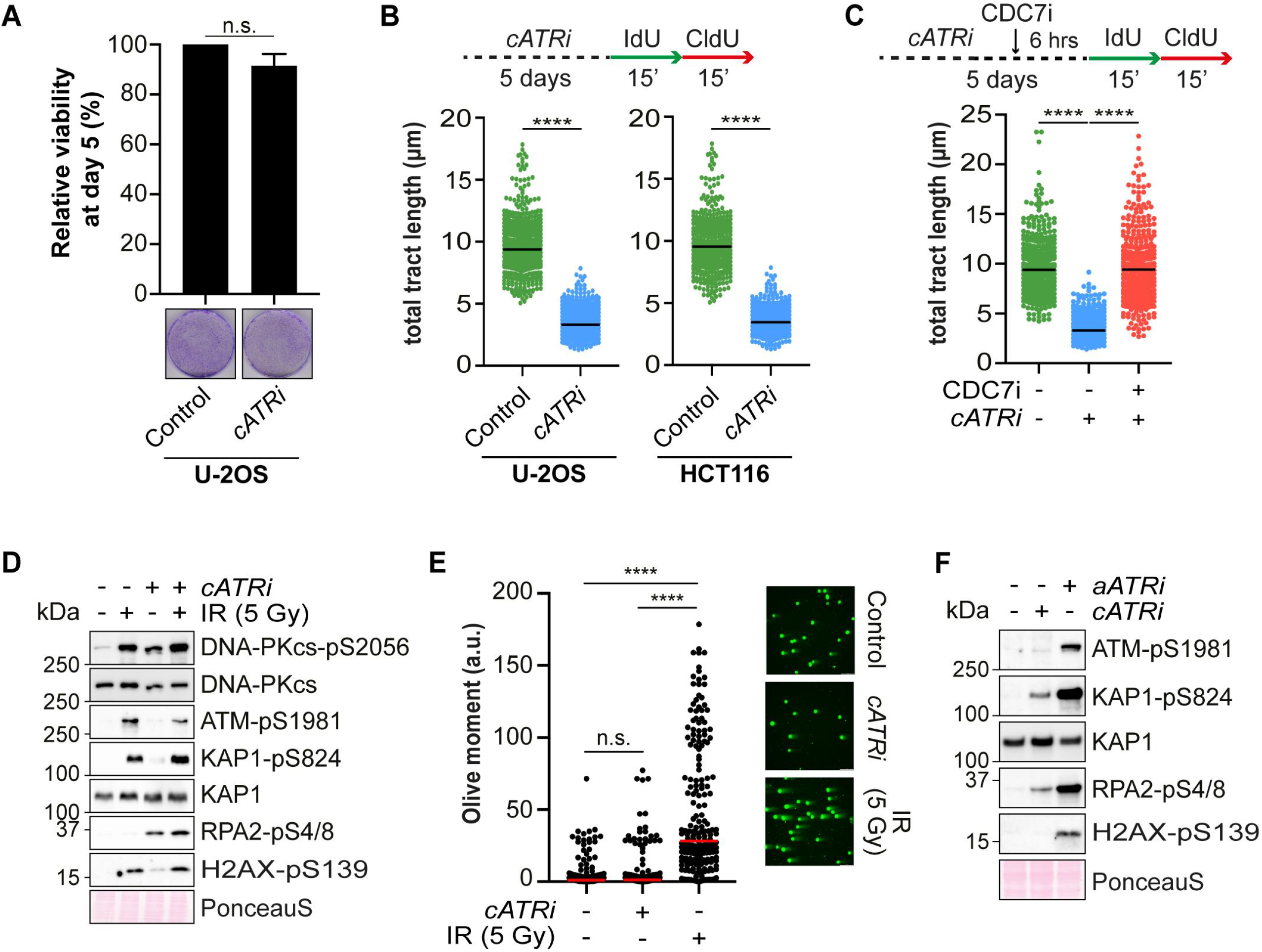
*cATRi* causes fork slowing without promoting extensive genomic instability. (A) U-2OS cells were treated for five days with 0.4µM AZD6738. At day five, cellular viability was assessed by Trypan blue staining. One plate was stained with Crystal violet as a representative image. Plotted results are the mean of three independent experiments ±SD. n.s. not significant (B) DNA fiber analysis of U-2OS and HCT116 cells grown as in (A). At least 400 individual fibers for each condition were scored. Dot plot and median of total tract length are shown (n=3). **** p<0.001 unpaired two-tailed Student’s *t*-test (C) U-2OS cells were treated as in (A) except that, on day 5, cells were additionally treated or mock with 10µM XL413 (CDC7i) for 6 hours. At least 350 individual fibers for each condition were scored. Dot plot and median of total tract length are shown (n=2). **** p<0.001 unpaired two-tailed Student’s *t*-test. (D) Western blot analysis of U-2OS cells irradiated (1 hour after 5 Gy) or treated with *cATRi*. (E) Neutral Comet assay of U-2OS cells treated as in (D). At least 200 individual nuclei for each condition were scored. Dot plot and median of olive moment are shown (n=2). n.s. not significant, **** p<0.001 One-way Anova test. (F) Western blot analysis of U-2OS cells treated with *cATRi* or *aATRi* (AZD6738, 5 µM for 8 hours).

### *cATRi* triggers fork reversal

Because forks are progressing at a much lower rate upon *cATRi*, we hypothesized that they might be stabilized through fork reversal (Berti et al., 2020a). We directly visualized *in vivo* fork architecture using psoralen crosslinking followed by Electron Microscopy (EM) in U-2OS cells following *cATRi* (Zellweger and Lopes, 2017) (Figure 2A). Consistent with our hypothesis, analysis of fork architecture upon *cATRi* revealed an approximately 4-fold increase in the frequency of reversed forks relative to control cells (Figure 2B).

**Figure 2.**
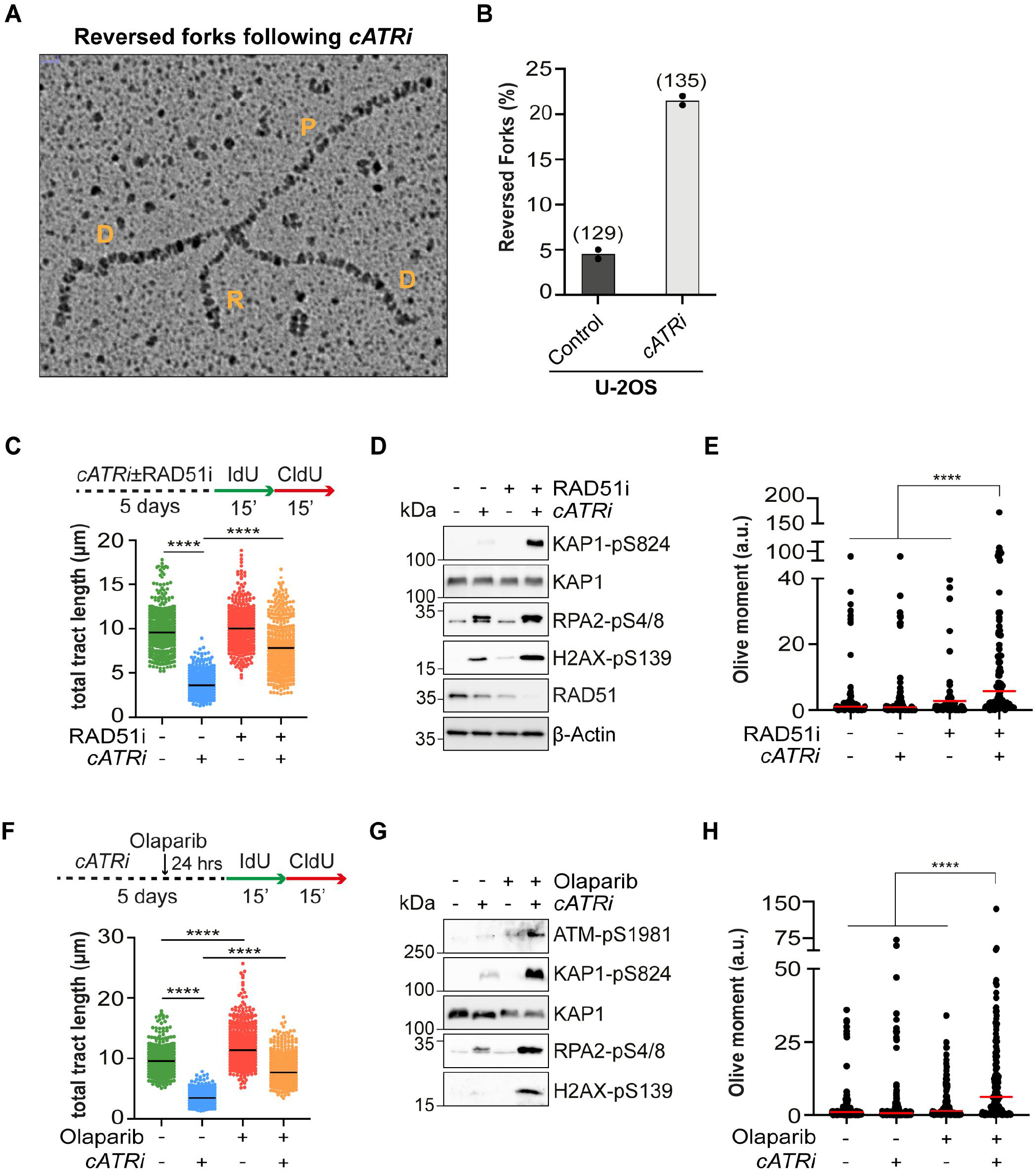
*cATRi* leads to frequent fork reversal. (A) Electron micrograph representing a reversed replication fork from U-2OS cells treated for five days with *cATRi*. P, parental strand; D, daughter strand; R, regressed arm. The arrow points to the regressed arm of the fork (B) Graph-bar showing the mean of reversed replication forks frequency following the indicated treatments from two independent EM experiments. In brackets, the total number of molecules analyzed is indicated. (C) DNA fiber analysis of U-2OS cells treated for five days with DMSO, 0.4µM AZD6738, 10µM B02 and their combination. At least 350 individual fibers for each condition were scored. Dot plot and median of total tract length are shown (n=3). **** p<0.001 One-way Anova test. (D) Western blot analysis of U-2OS cells grown as in (C). (E) Neutral Comet assay analysis of U-2OS cells treated as in (C). At least 100 individual nuclei for each condition were scored. Dot plot and median of olive moment are shown (n=2). n.s. not significant, **** p<0.001 One-way Anova test. (F) DNA Fiber analysis of U-2OS cells treated for four days with DMSO or 0.4µM AZD6738 and for additional 24 hours with ATRi with and without Olaparib (10µM). At least 400 individual fibers for each condition were scored. Dot plot and median of total tract length are shown (n=3). **** p<0.001 One-way Anova test. (G) Western blot analysis of U-2OS cells grown as in (F). (H) Neutral Comet assay analysis of U-2OS cells treated as in (F). At least 100 individual nuclei for each condition were scored. Dot plot and median of olive moment are shown (n=2). **** p<0.001 One-way Anova test.

Drug-induced fork slowing and reversal was shown to require the RAD51 remodeling activity (Berti et al., 2020b; Bugreev et al., 2011; Zellweger et al., 2015). To gain mechanistic insights into the process of altered fork dynamics following *cATRi*, we tested whether *cATRi*-induced fork slowing also depends on RAD51. We employed the B02 (RAD51i) small molecule, which disrupts RAD51 affinity for ssDNA (Huang et al., 2012). We combined B02 with *cATRi* treatment and tested fork speed by fiber assay (Figure 2C). Consistent with previous studies showing a role for RAD51 in mediating fork slowing and reversal in response to DNA damage (Berti et al., 2020b; Zellweger et al., 2015), B02 significantly rescued fork slowing triggered by *cATRi* (Figure 2C). Western blot analysis showed that RAD51 inhibition coupled with *cATRi* led to a drastic decrease in the levels of RAD51 protein (Figures 2D, S1A), which was also accompanied by a significant increase in the phosphorylation status of KAP1, RPA2, and H2AX (Figure 2D). The reduction in RAD51 levels and increased DSB signaling was prominent after five days of B02 and ATRi co-treatment, but only mild after a one-day co-treatment (Figure S1B). We also detected a substantial increase in double-strand break formation upon 5 days of RAD51i and *cATRi* co-treatment (Figure 2E), suggesting that failure to slowdown fork progression following *cATRi* generates fork breakage. Consistent with a role for RAD51 in limiting genome instability following *cATRi*, we observed a significant reduction in cellular viability in U-2OS and MDA-MB-231 cells after *cATRi*/RAD51i co-treatment (Figure S1C), and in HCT116-TetOn-shATR cells after doxycycline/RAD51i treatment (Figures S1C, D). Importantly, because RAD51i treatment did not trigger detectable ATR activation at the dose tested (Figure S1E), we do not attribute the *cATRi*/RAD51i toxicity to a loss of CHK1 activation.

Since PARP plays a pivotal role in stabilizing reversed forks (Berti et al., 2013), we next sought to investigate the ability of the PARPi (Olaparib) to accelerate fork speed following *cATRi*, with the expectation that abrogation of fork reversal via PARPi would restore fork speed (Figure 2F). Consistent with previous reports (Berti et al., 2013; Ray Chaudhuri et al., 2012), Olaparib treatment largely suppressed *cATRi*-induced fork slowing (Figure 2F). Similar results were also obtained in the HCT116 cell line (Figure S1F). Because suppression of fork speed by RAD51i led to the accumulation of DNA breaks upon *cATRi* (Figures 2D, E), we asked whether suppression of fork speed by PARPi would have a similar effect. Analysis of chromosome integrity by western blot analysis of DSB signaling markers and Neutral comet assay revealed that PARPi-mediated suppression of fork speed is accompanied by fork breakage (Figures 2G, H).

### Suppression of fork slowing by PARP inhibitors correlates with *cATRi*/PARPi toxicity

Recently, it has been proposed that acceleration of fork speed by PARPi drives DNA damage and genomic instability (Maya-Mendoza et al., 2018). To verify whether acceleration of fork speed by PARPi following *cATRi* correlates with impaired cellular proliferation, we employed distinct FDA-approved PARP inhibitors, Veliparib and Talazoparib, which have different potency and affect fork speed with different magnitude (Genois et al., 2021; Murai et al., 2013, 2014). Consistent with the results obtained with Olaparib (Figure 2F), we also observed suppression of *cATRi*-induced fork slowing with both Veliparib and Talazoparib (Figure 3A). The extent of fork speeding was correlated with the potency of the PARPi used, being higher for Talazoparib and lower with Veliparib (Genois et al., 2021; Murai et al., 2013, 2014). In line with the idea that fork reversal and fork slowing following *cATRi* is a key adaptive response protecting genome integrity, fork speed suppression by PARP inhibition resulted in a marked toxicity over a 5-day ATRi/PARPi co-treatment in U-2OS and HCT116 cells (Figures 3B, C). Importantly, the degree of sensitization also correlated with the extent of fork acceleration by the different PARPi (Figures 3B, C). In agreement with the idea that *cATRi*/PARPi toxicity derives from fork speed deregulation, rather than from loss of CHK1 activation, we could not detect any evidence of increased CHK1 phosphorylation with the concentration of Olaparib used (Figure 3D).

**Figure 3.**
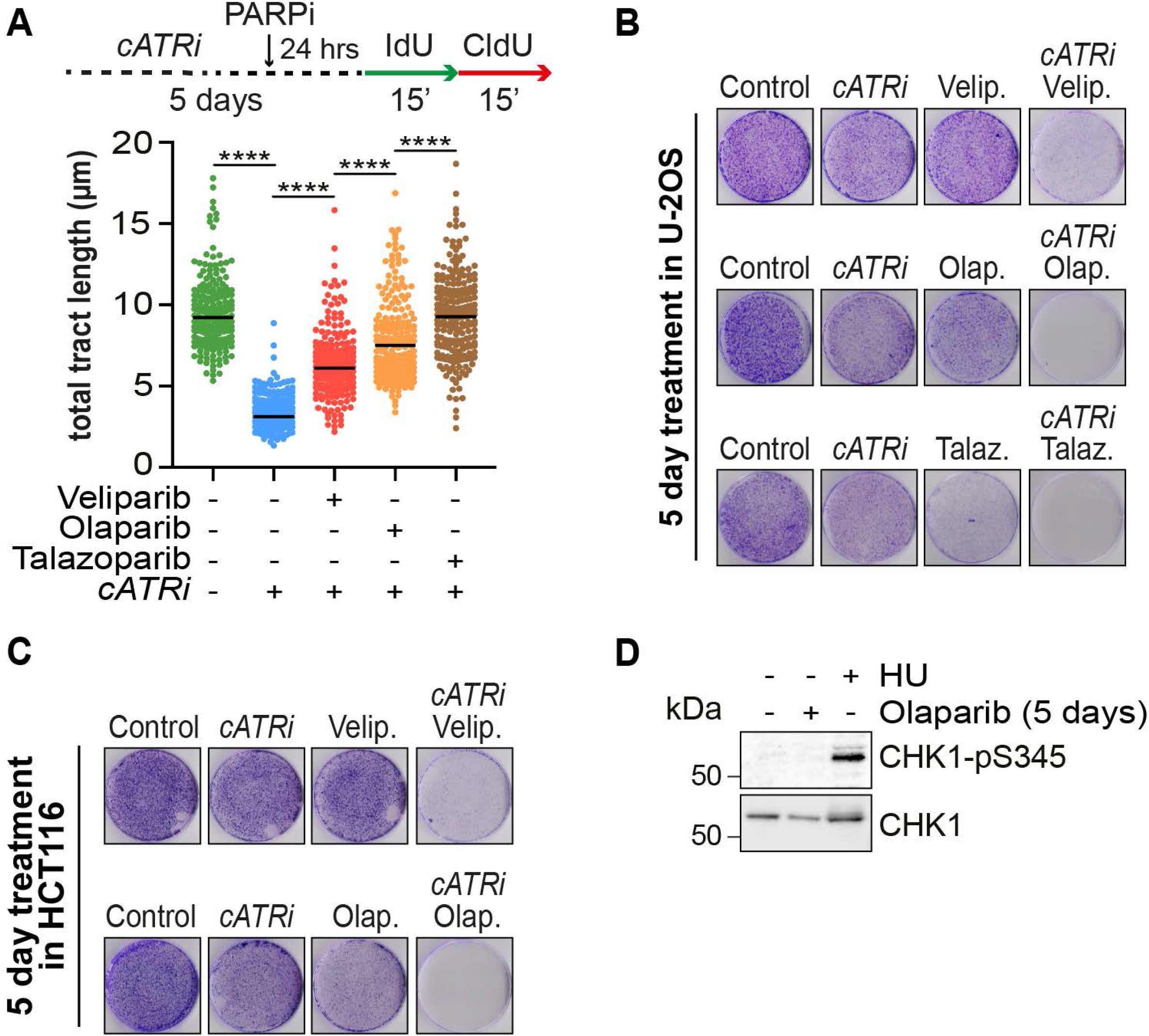
Fork acceleration by PARPi correlates with *cATRi*/PARPi toxicity. (A) DNA Fiber analysis of U-2OS cells treated as in Figure 2F, but with Veliparib (10µM), Olaparib (10µM) or Talazoparib (10µM). At least 200 individual fibers for each condition were scored. Dot plot and median of total tract length are shown (n=2). **** p<0.001 One-way Anova test. (B) U-2OS cells were treated for five days with the indicated combination of ATRi (AZD6738, 0.4µM), Veliparib (2.5 µM), Olaparib (2µM), and Talazoparib (20nM). On day five, cells were fixed and stained with Crystal Violet. (C) HCT116 cells were treated as in (B). (D) Western blot analysis of U-2OS cells treated for five days with Olaparib (2µM) or 2 hours with HU (1mM).

### Depletion of HR factors is dispensable for *cATRi*-induced PARPi hypersensitivity\

Recently, we have shown that *cATRi* affects the abundance of proteins involved in DNA end resection through the inhibition of E2F transcription (Dibitetto et al., 2020). For this reason, we asked whether impaired DNA end resection would cause the accumulation of reversed forks and slowed fork movement following *cATRi*. We employed two ovarian cancer cell lines, A2780 and SK-OV-3, where *cATRi* has distinct effects on HR factor abundance (Figure 4A). While *cATRi* resulted in drastic loss of the resection factors BRCA1, CTIP, and BLM in A2780 cells (Dibitetto et al., 2018, 2020), the abundance of resection factors in SK-OV-3 cells was completely refractory to the *cATRi* treatment (Figure 4A). Next, we analyzed fork dynamics in A2780 and SK-OV-3 cells following *cATRi* by DNA fiber assay. Surprisingly, analysis of fork speed revealed a similar reduction in the fork elongation rate in both A2780 and SK-OV-3 cells upon *cATRi* relative to the control (Figure 4B). These results indicate that loss of DNA end resection factors following *cATRi* plays a minor role in fork slowing and reversal. Consistent with this idea, we observed that, in SK-OV-3 cells, *cATRi*/PARPi still caused a drastic loss of viability (Figure 4C), which in this case we attributed to the loss of fork stabilization and not to the reduced abundance of HR factors and reduced resection capacity. Consistent with the strict PARP requirement for the slowing of forks following *cATRi*, Olaparib treatment largely suppressed fork speed following *cATRi* also in SK-OV-3 cells (Figure 4D), reinforcing the importance of PARP-mediated stabilization of reversed forks for preventing the deleterious consequences of *cATRi* for cell proliferation.

**Figure 4.**
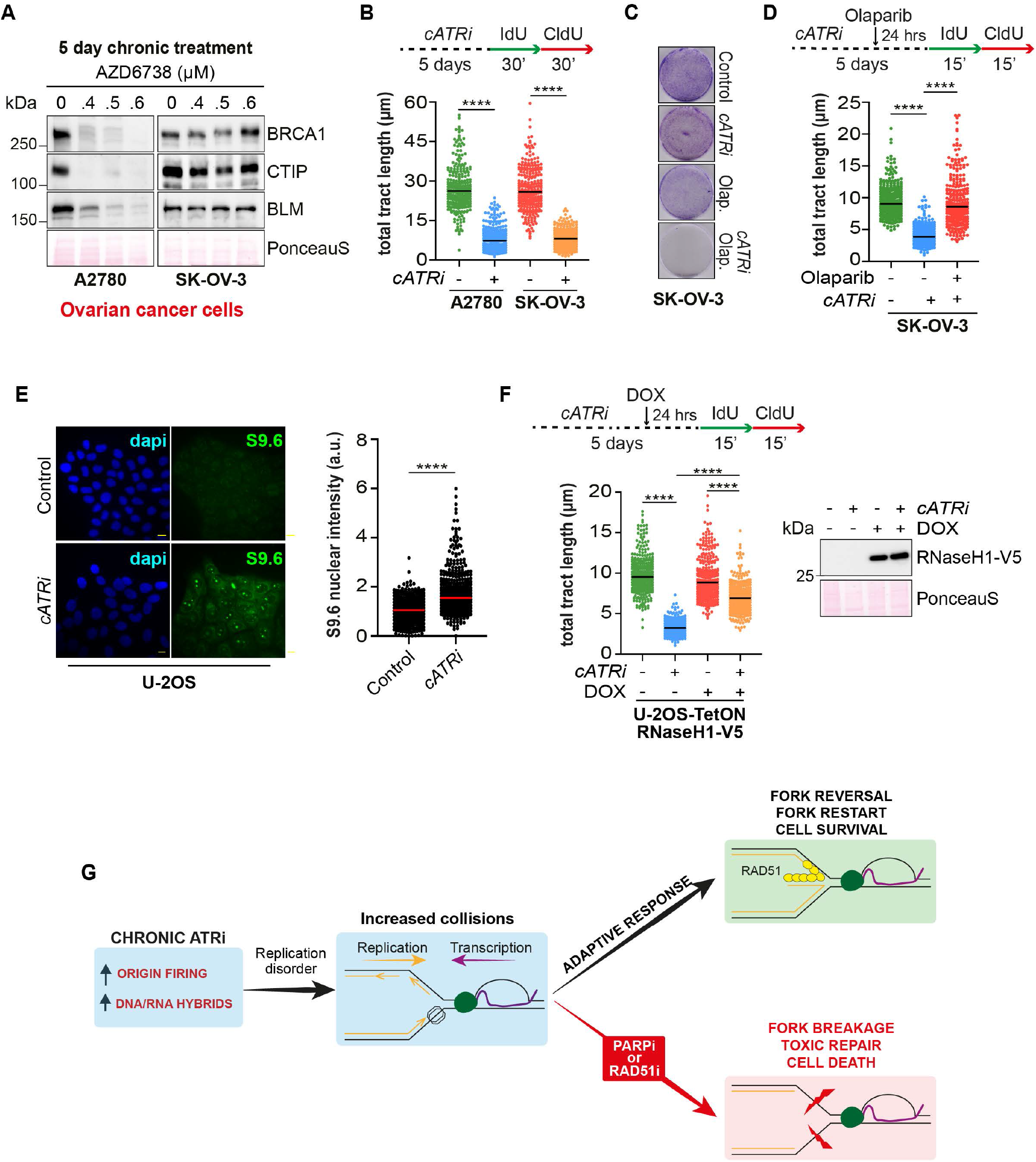
*cATRi*-mediated fork slowing does not depend on HR factor depletion but is associated with the accumulation of DNA-RNA hybrids. (A) Western blot analysis of A2780 and SK-OV-3 cells treated for five days with the indicated concentrations of AZD6738. (B) DNA Fiber analysis of A2780 and SK-OV-3 cells grown as in Figure 1B. At least 240 individual fibers for each condition were scored. Dot plot and median of total tract length are shown (n=2). **** p<0.001 One-way Anova test. (C) SK-OV-3 cells were treated for five days with the indicated combination of ATRi (AZD6738, 0.4µM) and Olaparib (2µM). On day five cells were fixed and stained with Crystal Violet. (D) DNA Fiber analysis of SK-OV-3 cells treated as in Figure 2F. At least 275 individual fibers for each condition were scored. Dot plot and median of total tract length are shown (n=3). **** p<0.001 One-way Anova test. (E) S9.6 immunofluorescence analysis of U-2OS cells treated or not with *cATRi*. 450 individual cells for each condition were scored. Dot plot is shown and the median of S9.6 nuclear intensity is expressed as arbitrary units (n=3). **** p<0.001 Two-tailed t test. Scale bar is 10µm (F) DNA Fiber analysis of U-2OS-TetON-RNaseH1-V5 cells treated for four days with *cATRi* and additional 24 hours with ATRi with and without doxycycline. At least 300 individual fibers for each condition were scored. Dot plot and median of total tract length are shown (n=3). **** p<0.001 One-way Anova test. (G) Model for the *cATRi*-induced fork slowing and reversal adaptive response.

Next we wanted to better define the molecular events associated with the slowing of active forks following *cATRi*. Recently, ATR signaling has been associated with the suppression of genomic R-loops (Barroso et al., 2019; Hodroj et al., 2017; Matos et al., 2020). Because transcription/replication collisions can trigger the formation of reversed forks and a subsequent reduction in fork speed (Chappidi et al., 2020), we asked whether accumulation of DNA-RNA hybrids may be the cause of the *cATRi*-induced fork slowing. Consistent with the recent literature, we found that *cATRi* significantly increased S9.6 nuclear signal in U-2OS cells, which indicates a higher level of DNA-RNA hybrids in the genome (Figure 4E). Next, we asked if induced processing of DNA-RNA hybrids could accelerate fork speed following *cATRi*. We ectopically expressed the RNaseH1 enzyme in U-2OS cells and measured fork elongation rate by DNA fiber assay. Surprisingly, RNaseH1 overexpression significantly rescued fork speed following *cATRi* (Figure 4F), indicating that the accumulation of DNA-RNA hybrids plays an important role in slowing fork movement following *cATRi*.

## DISCUSSION

ATR inhibitors (ATRi) are currently used in multiple pre-clinical trials for anti-cancer therapy, and a range of studies using cell lines have demonstrated the ability of ATRi to promote a remarkable sensitization of cancer cells to replication poisoning agents. Mechanistically, most of our understanding of how ATRi sensitize cells to these drugs derives from cell line-based experiments using acute inhibition of ATR. However, preclinical trials need to use chronic and low concentrations of ATRi since efficient inhibition of ATR results in documented organismal toxicity (Barnieh et al., 2021). Not surprisingly, most preclinical trials have opted to use a combination of non-toxic doses of ATRi with replication stress-inducing agents (Barnieh et al., 2021). Therefore, the acute and high dose treatments commonly used for experiments with cell lines are unlikely to reflect the therapeutic context in the clinic. Here we investigated the effects of *cATRi* on fork dynamics and cellular proliferation during unperturbed DNA replication. We observed that *cATRi* triggers fork reversal, which we propose is part of an adaptive cellular response that is crucial to maintain replisome stability and ensure cell survival.

A role for ATR in promoting fork reversal was recently demonstrated upon acute ATR inhibition and in the presence of DNA damaging agents, with less reversed forks accumulating following ATRi (Mutreja et al., 2018). Our data revealing an opposite effect for *cATRi* are surprising and likely reflect a distinctive form of replication stress triggered by *cATRi*, which leads to extensive fork reversal even upon impaired ATR activity. These findings further emphasize the importance of considering the ATRi delivery regime for understanding the impact of ATRi on genome metabolism, particularly in the context of cancer therapy. Noteworthy, an ATR’s anti-remodeling function, via ATR-dependent phosphorylation of the SMARCAL1 translocase, was previously proposed based on experiments using acute ATRi (Couch et al., 2013), however more work is needed to determine if reversed forks actually accumulate once this signaling event is impaired.

The mechanism and cause of fork reversal upon *cATRi* remain unclear. The rather frequent events of fork reversal observed following *cATRi* indicate that the equilibrium between fork regression and restoration is shifted towards the regressed state possibly due to the impairment of fork restart or lesion-bypass mechanisms. Indeed, *cATRi* may affect PRIMPOL stabilization and PRIMPOL-mediated repriming (Bai et al., 2020; Genois et al., 2021; Quinet et al., 2020) and could also affect the DNA2/WRN-mediated restart pathway, given the ATR requirement for WRN localization to stalled forks (Ammazzalorso et al., 2010).

Our findings consolidate an intricate connection between ATR signaling, fork reversal and fork speed. We propose that prolonged impairment of ATR signaling concomitantly elevates origin firing and DNA-RNA hybrids, increasing Transcription/Replication Collisions (TRCs) that underpin the formation of reversed forks (Figure 4C). It is worth mentioning that our data connecting ATR signaling, TRCs and fork reversal gains support from an earlier work in budding yeast, where the Mec1-Rad53 checkpoint axis was shown to control the tethering of transcriptional units to the nucleopore, limiting the topological stress in front of ongoing forks and their subsequent reversal (Bermejo et al., 2011). More recently, this model gained confirmation also in human cells, where ATR has been proposed to regulate mRNA processing factors and alleviate the topological stress associated with splicing deficiency (Teloni et al., 2019).

Overall, our findings showing that *cATRi* triggers an adaptive response characterized by fork reversal and slowing will better serve in the rational design of more effective and targeted anti-cancer therapies. Indeed, our work reveals that inhibition of proteins required for fork reversal, such as RAD51 and PARP, hypersensitize cells to *cATRi* by impairing such adaptive response. In agreement with recent data showing that PARPi may act through the acceleration of fork speed and the subsequent formation of ssDNA gaps (Maya-Mendoza et al., 2018; Panzarino et al., 2020), we do not observe significant double-strand break formation and CHK1 activation in PARPi-treated cells after 5 days. This finding strongly supports the idea that *cATRi* and PARPi synthetic lethality cannot be simplistically explained by the loss of PARP DNA repair functions or by the loss of ATR checkpoint functions. Instead, we now propose that the reasons behind the hypersensitization observed upon *cATRi* and PARPi combination are likely multifactorial and complex, and part of the cell sensitivity to ATRi/PARPi could reflect the inability to stabilize the replisome through fork reversal. For these reasons and given that deregulation of DNA replication is nearly ubiquitously in tumors, we propose that the combination of *cATRi* with drugs that prevent fork reversal, such as PARPi, will find broad applications in cancer therapy.

## ACKNOWLEDGEMENTS

We thank Fenghua Hu for the access to the Fluorescent Microscope. We thank Jumana Badar for troubleshooting the Neutral Comet assay technique. We thank Beatriz Almeida for technical assistance. We thank Carolline F.R. Ascençao for the TetON-shATR plasmid. We thank Xiaojing Ma for sharing SK-OV-3 cells. We thank all the members of the Smolka lab for helpful discussions. D.D. is supported by a postdoctoral fellowship from Fleming Research Foundation. This work was supported by a NIH grants RO1-GM097272 and R35-GM141159 to M.B.S.

## AUTHOR CONTRIBUTIONS

D.D. and M.B.S. conceptualized the project. D.D. and E.J.S. performed experiments. A.S. and M.L. conducted EM analysis of reversed forks. D.D. and M.B.S. wrote the manuscript. All authors contributed to revising and editing the manuscript.

## STAR METHODS

## LEAD CONTACT AND MATERIALS AVAILABILITY

Further information and request for resources and reagents should be directed to and will be fulfilled by the Lead Contact, Marcus Smolka. All unique/stable reagents generated in this study are available from the Lead Contact without restriction.

## DATA AND CODE AVAILABILITY

Western blot raw data are available upon request.

## EXPERIMENTAL MODEL AND SUBJECT DETAILS

Source of cell lines used in the study is reported in the Key Resources Tables.

### Cell culture and cell lines

U-2OS, HCT116, MDA-MB-231, A2780, and 293T were grown at 37°C in humidified chambers in Dulbecco’s modified Eagle medium (DMEM) supplemented with 10% bovine calf serum (BCS) supplemented with 1% non-essential aminoacids and penicillin and streptomycin (100 U/ml). SK-OV-3 cells were grown in RPMI 1640 medium supplemented with 1% non-essential aminoacids and penicillin and streptomycin (100 U/ml). HCT116-TetON-shATR cells were maintained in DMEM supplemented with 10% bovine calf serum (BCS), 1% non-essential aminoacids, penicillin and streptomycin (100 U/ml) and puromycin (1 µg/ml). ATR knockdown was induced for five days with doxycycline (1 µg/ml). U-2OS-TetON-RNaseH1-V5 cells were maintained in DMEM supplemented with 10% bovine calf serum (BCS), 1% non-essential aminoacids, penicillin and streptomycin (100 U/ml) and G418 (500 µg/ml). RnasH1-V5 overexpression was induced with doxycycline (1 µg/ml).

## METHOD DETAILS

### Oligos and plasmids

The pCW57-TetON-RNaseH1-V5 plasmid was generated by Gibson Assembly with the following primers: RNH1_ins_F 3’-AATTCGTCGACACCGGTTCCGG AATGCCCAAGAAGAAGAGG-5’, RNH1_ins_R 3’-GTACAACGCGTCTGCAGCCTTCA CGTAGAATCGAGACC-5’, RNH1_int_F 3’-CCTGGAA TGAGTGCAGAGC-5’, RNH1_int_R 3’-TGCATGAATTTCCGCTCTTTGG-5’. pCW57-MCS1-P2A-MCS2 (Neo) was a gift from Adam Karpf (Addgene plasmid #89180; http://n2t.net/addgene:89180; RRID: Addgene_89180). ppyCAG_RNaseH1_WT was a gift from Xiang-Dong Fu (Addgene plasmid #111906; http://n2t.net/addgebe:111906; RRID:Addgene_111906). The pLKO-TetON-shATR plasmid was generated by inserting the following annealed oligos: shRNA-ATR-F1 3’-CCGGAAGGACATGTGCATTACCTTACTCGAGTAAGGTAATGCACATGTCC TTTTTTTG-5’, shRNA-ATR-R1 3’-AATTCAAAAAAAGGACATGTGCATTACCTTACTCGAGTAAGGTAATGCACATGTCCTT-5’. Tet-pLKO-puro was a gift from Dmitri Wiederschain (Addgene plasmid #21915; http://n2t.net/addgene:21915; RRID: Addgene_21915).

### Drugs and reagents

The following chemical reagents were used throughout the study: AZD6738 (Selleck Chemicals; #S7693), XL413 (Selleck Chemicals; #S7547), B02 (Selleck Chemicals; #S8434), Olaparib (Selleck Chemicals; #S1060), Veliparib (Selleck Chemicals; #S1004), Talazoparib (Selleck Chemicals; #S7048), 5-Iodo-2’-deoxyuridine (IdU) (Millipore Sigma; #I7125), 5-Chloro-2’-deoxyuridine (CldU) (Millipore Sigma; #C6891), Hydroxyurea (Tokyo Chemical Industry; #H0310), Doxycycline (Millipore Sigma; #D1822).

### Western blot analysis

Cells were lysed in ice-cold modified RIPA buffer (50mM Tris-HCl, pH 7.5, 150mM NaCl, 1% Tergitol, 0.25% sodium deoxycholate, 5mM EDTA supplemented with complete EDTA-free protease inhibitor cocktail (Roche), 1mM PMSF and 5mM NaF) for 30’ at 4°C. Lysates were then sonicated and cleared by centrifugation. Protein extracts were quantified with Bradford (Bio-Rad), and 20 µg of proteins was resolved by SDS-PAGE. Proteins were transferred to polyvinylidene difluoride (PVDF) membranes and chemiluminescent signal was acquired with a Chemidoc Imaging System (Bio-Rad). Images were processed with the ImageLab software (Bio-Rad).

### Antibodies

The following primary antibodies were used: anti-β-Actin (Thermo Fisher Scientific; #MA1-140), anti-pDNA-PKcs-S2056 (Thermo Fisher Scientific; #PA5-78130), anti-DNA-PKcs (Bethyl Laboratories; #A300-516A), anti-pRPA2-S4/8 (Bethyl Laboratories; #A300-245A), anti-RAD51 (Merck Millipore; #PC130), anti-pKAP1-S824 (Bethyl Laboratories; #A300-767A), anti-pATM-S1981 (Cell Signaling Technology; #13050S), anti-KAP1 (Bethyl Laboratories; #A300-274A), anti-pCHK1-S345 (Cell Signaling Technology; #2341), anti-CHK1 (Santa Cruz Biotechnology; #sc-8408), anti-pH2AX-S139 (Bethyl Laboratories; A300-081A), anti-BRCA1 (Cell Signaling Technology; #9010), anti-CTIP (Bethyl Laboratories; #A300-488A), anti-BLM (Bethyl Laboratories; #A300-110A), anti-V5 (Thermo Fisher Scientific; #R960-25). The secondary antibodies used were Goat anti-Mouse IgG-HRP (Thermo Fisher Scientific; #NA931V) and Donkey anti-Rabbit IgG-HRP (Thermo Fisher Scientific; #NA934V).

### Neutral Comet assay

DNA double-strand breaks were detected by Neutral Comet assay using a protocol from Trevigen. U-2OS cells treated with the indicated drugs were mixed with molten LMAgarose at a ratio of 1:10 and then immediately pipetted onto Comet slides (Trevigen). Slides were soaked in cold Lysis Solution (Trevigen) for 1 hour. Slides were then soaked for 30’ in Neutral Electrophoresis Buffer (Tris Base, Sodium Acetate, pH 9.0), then placed in the electrophoresis apparatus with a constant 20V setting for 30’. Slides were immersed in DNA precipitation solution (Ammonium Acetate, 95% Ethanol) for 30’ followed by 30’ in 70% Ethanol. Slides were dried, stained with SYBR Safe and imaged using a Leica DMi8 inverted fluorescent microscope with a 40x objective. For each condition, an average of 100-200 cells were scored using the OpenComet software. Statistical analysis of Olive moment was performed using Prism (GraphPad Software) and was determined with the one-way ANOVA test.

### DNA Fiber assay

We analyzed DNA fiber length as previously described (Quinet et al., 2017). In brief, exponentially growing cells were first labeled with 25 µM IdU for 15’ followed by 250 µM CldU 15’ incubation. Cells were then harvested by centrifugation, washed with PBS, and approximately 2,000 cells were spotted onto glass slides. Cells were mixed on the slide with lysis buffer (200mM Tris-HCl pH 7.5, 50mM EDTA, 0.5% SDS) for 3’. After this incubation step, slides were tilted at 30° to allow uniform fiber spreading. Slides were air-dried for 5’, fixed at RT for 10’ in Methanol-Acetic Acid (3:1), and stored at 4°C overnight. The day after, slides were denatured 1 hour in 2.5M HCl, quickly washed in PBS, and blocked for 1 hour in 10% PBS-BSA. Newly replicated DNA was stained for 2 hours with primary antibodies anti-IdU (mouse, BD Biosciences) and anti-CldU (rat, Abcam). Slides were then washed three times in PBS followed by 1 hour with the secondary antibodies anti-mouse Alexa Fluor 488 and anti-rat Alexa Fluor 594. Slides were washed three times in PBS, mounted with Fluoromount G (Thermo Fisher Scientific), and sealed with nail polish. Images were acquired using a Leica DMi8 inverted fluorescent microscope with a 63x objective. The images were processed with the ImageJ software. Statistical analysis of fiber length was performed using Prism (GraphPad Software). Statistical significance was determined by Student *t*-test if comparing two conditions or one-way ANOVA test for multiple comparisons.

### Immunofluorescence analysis of DNA-RNA hybrids

U-2OS cells were seeded on coverslips and subjected to the indicated treatments. On the last day, cells were washed in PBS and fixed with ice-cold methanol at -20°C for 20’. Cells were then permeabilized for 5’ and blocked for 20’ with 10%PBS-BSA at RT. For DNA-RNA hybrids staining, coverslips were incubated O/N at 4°C with a monoclonal S9.6 antibody (Kerafast, 1:100) diluted in 3%PBS-BSA. The day after, coverslips were washed three times 5’ with PBS followed by 1-hour incubation at RT with a secondary antibody (Alexafluor 488, 1:600). Coverslips were washed three times with PBS and then mounted with DAPI on microscope slides. Images were acquired using a Leica DMi8 inverted fluorescent microscope with a 63x objective. For measuring the nuclear S9.6 intensity, the images were processed with the ImageJ software. Statistical analysis was performed using Prism (GraphPad Software). Statistical significance was determined by two tailed *t*-test.

### Electron Microscopy (EM)

Asynchronous sub confluent U-2OS cells were treated for five days with 0.4uM AZD6738 or DMSO. Cells were collected, resuspended in ice-cold PBS and crosslinked with 4,5′, 8-trimethylpsoralen (10 μg/ml final concentration), followed by irradiation pulses with UV 365 nm monochromatic light (UV Stratalinker 1800; Agilent Technologies). For DNA extraction, cells were lysed (1.28 M sucrose, 40 mM Tris-HCl [pH 7.5], 20 mM MgCl2, and 4% Triton X-100; Qiagen) and digested (800 mM guanidine–HCl, 30 mM Tris-HCl [pH 8.0], 30 mM EDTA [pH 8.0], 5% Tween-20, and 0.5% Triton X-100) at 50 °C for 2 h in presence of 1 mg/ml proteinase K. The DNA was purified using chloroform/isoamylalcohol (24:1) and precipitated in 0.7 volume of isopropanol. Finally, the DNA was washed with 70% EtOH and resuspended in 200μl TE (Tris-EDTA) buffer. 100 U of restriction enzyme (PvuII high fidelity, New England Biolabs) were used to digest 6μg of mammalian genomic DNA for 5 h. RNase A (Sigma– Aldrich, R5503) to a final concentration of 250 ug/ml was added for the last 2 h of this incubation. The digested DNA was cleaned-up using the Silica Bead DNA Gel Extraction Kit (Thermo Fisher Scientific). The Benzyldimethylalkylammonium chloride (BAC) method was used to spread the DNA on the water surface and then load it on carbon-coated 400-mesh nickel grids (G2400N, Plano Gmbh). Subsequently, DNA was coated with platinum using a High Vacuum Evaporator BAF060 (Leica). The grids were scanned using a transmission electron microscope (Tecnai G2 Spirit; FEI; LaB6 filament; high tension ≤ 120 kV) and pictures were acquired with a side mount charge-coupled device camera (2600 × 4000 pixels; Orius 1000; Gatan, Inc.) and processed with Digital Micrograph Version 1.83.842 (Gatan, Inc.). For each experimental condition at least 65 replication fork molecules were analyzed with the ImageJ software.

### Viability assay

25,000-50,000 cells were seeded on 6 cm Petri dishes in the presence of the indicated drugs and grown for five days. At the end of the experiment, plates were washed with PBS, fixed for 30’ with 3.7% Formaldehyde, and stained with Crystal Violet.

### Statistical analysis

Statistical analysis was done with the one-way ANOVA test using Prism (GraphPad Software) when comparing three or more datasets. For comparison of two datasets, we determined statistical significance using the two-tailed t test. In all cases, n.s. indicates not significant, * (P<0.05), ** (P<0.01), *** (P<0.001), **** (P<0.0001). More details can be found in each figure legend.

## DATA AND CODE AVAILABILITY

All data are available by request.

---

**Figure S1.**
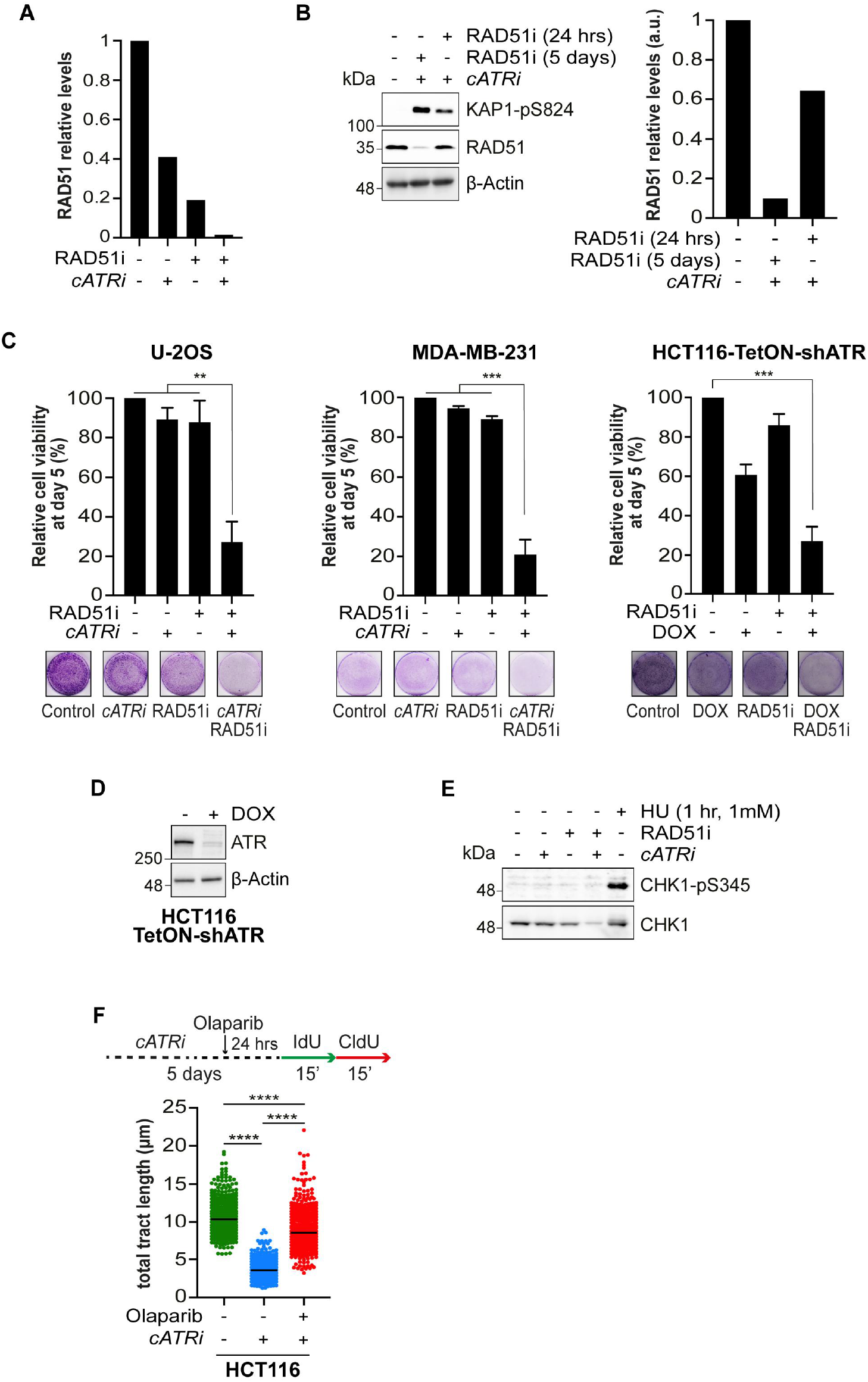
RAD51 suppresses genome instability following *cATRi*. (A) RAD51 quantification relative to the Western blot shown in Figure 2B. (B) Immunoblot analysis, and relative quantification, of U-2OS cells treated with the indicated combination of AZD6738 and B02. (C) U-2OS and MDA-MB-231 were treated with the indicated combination of ATRi (AZD6738, 0.4µM) and RAD51i (B02, 10µM) for 5 days. For HCT116-TetON-shATR, cells were treated with the indicated combination of doxycycline and RAD51i (B02, 10µM) for five days. On day five, cells were harvested, and cell viability was assessed by Trypan blue staining. A representative image of was obtained by crystal violet staining (see the STAR Methods). Plotted results are the mean of two independent experiments ±SD. ** p<0.1, *** p<0.01 One-way Anova test. (D) Immunoblot analysis of HCT116-TetON-shATR cells treated for five days with or without doxycycline. (E) Immunoblot analysis of U-2OS cells treated with the indicated combination of AZD6738 (0.4µM), B02 (10µM), and HU (1mM). (F) HCT116 cells were treated as described in Figure 2D. The DNA fiber protocol was done following the conditions as in Figure 1B. 400 individual fibers for each condition were scored. Dot plot and median of total tract length are shown (n=2). **** p<0.001 One-way Anova test.

## REFERENCES

Ammazzalorso, F., Pirzio, L.M., Bignami, M., Franchitto, A., and Pichierri, P. (2010). ATR and ATM differently regulate WRN to prevent DSBs at stalled replication forks and promote replication fork recovery. EMBO J. 29, 3156–3169.

Bai, G., Kermi, C., Stoy, H., Schiltz, C.J., Bacal, J., Zaino, A.M., Hadden, M.K., Eichman, B.F., Lopes, M., and Cimprich, K.A. (2020). HLTF Promotes Fork Reversal, Limiting Replication Stress Resistance and Preventing Multiple Mechanisms of Unrestrained DNA Synthesis. Mol. Cell 78(6), 1237–1251. e7.

Barnieh, F.M., Loadman, P.M., and Falconer, R.A. (2021). Progress towards a clinically-successful ATR inhibitor for cancer therapy. Curr. Res. Pharmacol. Drug Discov. 2, 100017.

Barroso, S., Herrera-Moyano, E., Muñoz, S., García-Rubio, M., Gómez-González, B., and Aguilera, A. (2019). The DNA damage response acts as a safeguard against harmful DNA– RNA hybrids of different origins. EMBO Rep. 20(9), e47250.

Bermejo, R., Capra, T., Jossen, R., Colosio, A., Frattini, C., Carotenuto, W., Cocito, A., Doksani, Y., Klein, H., Gómez-González, B., et al. (2011). The replication checkpoint protects fork stability by releasing transcribed genes from nuclear pores. Cell 146(2), 233–246.

Berti, M., Chaudhuri, A.R., Thangavel, S., Gomathinayagam, S., Kenig, S., Vujanovic, M., Odreman, F., Glatter, T., Graziano, S., Mendoza-Maldonado, R., et al. (2013). Human RECQ1 promotes restart of replication forks reversed by DNA topoisomerase I inhibition. Nat. Struct. Mol. Biol. 20(3), 347–354.

Berti, M., Cortez, D., and Lopes, M. (2020a). The plasticity of DNA replication forks in response to clinically relevant genotoxic stress. Nat. Rev. Mol. Cell Biol. 21(10), 633–651.

Berti, M., Teloni, F., Mijic, S., Ursich, S., Fuchs, J., Palumbieri, M.D., Krietsch, J., Schmid, J.A., Garcin, E.B., Gon, S., et al. (2020b). Sequential role of RAD51 paralog complexes in replication fork remodeling and restart. Nat. Commun. 11(1), 3531.

Bugreev, D. V., Rossi, M.J., and Mazin, A. V. (2011). Cooperation of RAD51 and RAD54 in regression of a model replication fork. Nucleic Acids Res. 39(6), 2153–2164.

Buisson, R., Boisvert, J.L., Benes, C.H., and Zou, L. (2015). Distinct but Concerted Roles of ATR, DNA-PK, and Chk1 in Countering Replication Stress during S Phase. Mol. Cell 59(6), 1011–1024.

Byun, T.S., Pacek, M., Yee, M.C., Walter, J.C., and Cimprich, K.A. (2005). Functional uncoupling of MCM helicase and DNA polymerase activities activates the ATR-dependent checkpoint. Genes Dev. 19(9), 1040–1052.

Chappidi, N., Nascakova, Z., Boleslavska, B., Zellweger, R., Isik, E., Andrs, M., Menon, S., Dobrovolna, J., Balbo Pogliano, C., Matos, J., et al. (2020). Fork Cleavage-Religation Cycle and Active Transcription Mediate Replication Restart after Fork Stalling at Co-transcriptional R-Loops. Mol. Cell 77(3), 528–541.e8.

Charrier, J.D., Durrant, S.J., Golec, J.M.C., Kay, D.P., Knegtel, R.M.A., MacCormick, S., Mortimore, M., O’Donnell, M.E., Pinder, J.L., Reaper, P.M., et al. (2011). Discovery of Potent and Selective Inhibitors of Ataxia Telangiectasia Mutated and Rad3 Related (ATR) Protein Kinase as Potential Anticancer Agents. J. Med. Chem. 54(7), 2320–2330.

Checkley, S., Maccallum, L., Yates, J., Jasper, P., Luo, H., Tolsma, J., and Bendtsen, C. (2015). Bridging the gap between in vitro and in vivo: Dose and schedule predictions for the ATR inhibitor AZD6738. Sci. Rep. 5, 13545.

Costanzo, V., Shechter, D., Lupardus, P.J., Cimprich, K.A., Gottesman, M., and Gautier, J. (2003). An ATR- and Cdc7-dependent DNA damage checkpoint that inhibits initiation of DNA replication. Mol. Cell 11(1), 203–213.

Couch, F.B., Bansbach, C.E., Driscoll, R., Luzwick, J.W., Glick, G.G., Bétous, R., Carroll, C.M., Jung, S.Y., Qin, J., Cimprich, K.A., et al. (2013). ATR phosphorylates SMARCAL1 to prevent replication fork collapse. Genes Dev. 27(14), 1610–1623.

Dibitetto, D., La Monica, M., Ferrari, M., Marini, F., and Pellicioli, A. (2018). Formation and nucleolytic processing of Cas9-induced DNA breaks in human cells quantified by droplet digital PCR. DNA Repair (Amst). 68, 68–74.

Dibitetto, D., Sims, J.R., Ascencao, C.F.R., Feng, K., Kim, D., Oberly, S., Freire, R., and Smolka, M.B. (2020). Intrinsic ATR signaling shapes DNA end resection and suppresses toxic DNA-PKcs signaling. Nucleic Acids Res. Cancer 2(2), 1–14.

Foote, K.M., Nissink, J.W.M., McGuire, T., Turner, P., Guichard, S., Yates, J.W.T., Lau, A., Blades, K., Heathcote, D., Odedra, R., et al. (2018). Discovery and Characterization of AZD6738, a Potent Inhibitor of Ataxia Telangiectasia Mutated and Rad3 Related (ATR) Kinase with Application as an Anticancer Agent. J. Med. Chem. 61(22), 9889–9907.

Fròsina, G., Profumo, A., Marubbi, D., Marcello, D., Ravetti, J.L., and Daga, A. (2018). ATR kinase inhibitors NVP-BEZ235 and AZD6738 effectively penetrate the brain after systemic administration. Radiat. Oncol. 13(1), 76.

Genois, M.-M., Gagné, J.-P., Yasuhara, T., Jackson, J., Saxena, S., Langelier, M.-F., Ahel, I., Bedford, M.T., Pascal, J.M., Vindigni, A., et al. (2021). CARM1 regulates replication fork speed and stress response by stimulating PARP1. Mol. Cell 81(4), 784–800. e8.

Hodroj, D., Recolin, B., Serhal, K., Martinez, S., Tsanov, N., Abou Merhi, R., and Maiorano, D. (2017). An ATR -dependent function for the Ddx19 RNA helicase in nuclear R-loop metabolism. EMBO J. 36(9), 1182–1198.

Huang, F., Mazina, O.M., Zentner, I.J., Cocklin, S., and Mazin, A. V. (2012). Inhibition of homologous recombination in human cells by targeting RAD51 recombinase. J. Med. Chem. 55(7), 3011–3020.

Kiesel, B.F., Shogan, J.C., Rachid, M., Parise, R.A., Vendetti, F.P., Bakkenist, C.J., and Beumer, J.H. (2017). LC–MS/MS assay for the simultaneous quantitation of the ATM inhibitor AZ31 and the ATR inhibitor AZD6738 in mouse plasma. J. Pharm. Biomed. Anal. 138, 158–165.

Kim, D., Liu, Y., Oberly, S., Freire, R., and Smolka, M.B. (2018). ATR-mediated proteome remodeling is a major determinant of homologous recombination capacity in cancer cells. Nucleic Acids Res. 46(16), 8311–8325.

Kim, H., George, E., Ragland, R.L., Rafail, S., Zhang, R., Krepler, C., Morgan, M.A., Herlyn, M., Brown, E.J., and Simpkins, F. (2017). Targeting the ATR/CHK1 axis with PARP inhibition results in tumor regression in BRCA-mutant ovarian cancer models. Clin. Cancer Res. 23(12), 3097–3108.

Kim, H., Xu, H., George, E., Hallberg, D., Kumar, S., Jagannathan, V., Medvedev, S., Kinose, Y., Devins, K., Verma, P., et al. (2020). Combining PARP with ATR inhibition overcomes PARP inhibitor and platinum resistance in ovarian cancer models. Nat. Commun. 11(1), 3726.

Lanz, M.C., Dibitetto, D., and Smolka, M.B. (2019). DNA damage kinase signaling: checkpoint and repair at 30 years. EMBO J. 38(18), e101801.

Lecona, E., and Fernandez-Capetillo, O. (2018). Targeting ATR in cancer. Nat. Rev. Cancer 18(9), 586–595.

Lloyd, R.L., Wijnhoven, P.W.G., Ramos-Montoya, A., Wilson, Z., Illuzzi, G., Falenta, K., Jones, G.N., James, N., Chabbert, C.D., Stott, J., et al. (2020). Combined PARP and ATR inhibition potentiates genome instability and cell death in ATM-deficient cancer cells. Oncogene 39(25), 4869–4883.

Matos, D.A., Zhang, J.M., Ouyang, J., Nguyen, H.D., Genois, M.M., and Zou, L. (2020). ATR Protects the Genome against R Loops through a MUS81-Triggered Feedback Loop. Mol. Cell 77(3), 514–527. e4.

Maya-Mendoza, A., Moudry, P., Merchut-Maya, J.M., Lee, M., Strauss, R., and Bartek, J. (2018). High speed of fork progression induces DNA replication stress and genomic instability. Nature 559(7713), 279–284.

Moiseeva, T., Hood, B., Schamus, S., O’Connor, M.J., Conrads, T.P., and Bakkenist, C.J. (2017). ATR kinase inhibition induces unscheduled origin firing through a Cdc7-dependent association between GINS and And-1. Nat. Commun. 8(1), 1392.

Moiseeva, T.N., Yin, Y., Calderon, M.J., Qian, C., Schamus-Haynes, S., Sugitani, N., Osmanbeyoglu, H.U., Rothenberg, E., Watkins, S.C., and Bakkenist, C.J. (2019). An ATR and CHK1 kinase signaling mechanism that limits origin firing during unperturbed DNA replication. Proc. Natl. Acad. Sci. U. S. A. 116(27), 13374–13383.

Murai, J., Huang, S.N., Das, B.B., Renaud, A., Zhang, Y., Doroshow, J.H., Ji, J., Takeda, S., and Pommier, Y. (2013). Differential trapping of PARP1 and PARP2 by clinical Parp inhibitors. Cancer Res. 72(21), 5588–5599.

Murai, J., Zhang, Y., Morris, J., Ji, J., Takeda, S., Doroshow, J.H., and Pommier, Y. (2014). Rationale for poly(ADP-ribose) polymerase (PARP) inhibitors in combination therapy with camptothecins or temozolomide based on PARP trapping versus catalytic inhibition. J. Pharmacol. Exp. Ther. 349(3), 408–416.

Mutreja, K., Krietsch, J., Hess, J., Ursich, S., Berti, M., Roessler, F.K., Zellweger, R., Patra, M., Gasser, G., and Lopes, M. (2018). ATR-Mediated Global Fork Slowing and Reversal Assist Fork Traverse and Prevent Chromosomal Breakage at DNA Interstrand Cross-Links. Cell Rep. 24(10), 2629–2642. e5.

Panzarino, N.J., Krais, J.J., Cong, K., Peng, M., Mosqueda, M., Nayak, S.U., Bond, S.M., Calvo, J.A., Doshi, M.B., Bere, M., et al. (2020). Replication Gaps Underlie BRCA-deficiency and Therapy Response. Cancer Res. 81(5), 1388–1397.

Petermann, E., Woodcock, M., and Helleday, T. (2010). Chk1 promotes replication fork progression by controlling replication initiation. Proc. Natl. Acad. Sci. U. S. A. 107(37), 16090–16095.

Quinet, A., Carvajal-Maldonado, D., Lemacon, D., and Vindigni, A. (2017). DNA Fiber Analysis: Mind the Gap!. Methods Enzymol. 591, 55–82.

Quinet, A., Tirman, S., Jackson, J., Me, J., Sale, J.E., Vindigni, A., Quinet, A., Tirman, S., and Jackson, J. (2020). PRIMPOL-Mediated Adaptive Response Suppresses Replication Fork Reversal in BRCA-Deficient Cells. Mol. Cell 77(3), 1–14. e9.

Rainey, M.D., Bennett, D., O’Dea, R., Zanchetta, M.E., Voisin, M., Seoighe, C., and Santocanale, C. (2020). ATR Restrains DNA Synthesis and Mitotic Catastrophe in Response to CDC7 Inhibition. Cell Rep. 32(9), 108096.

Ray Chaudhuri, A., Hashimoto, Y., Herrador, R., Neelsen, K.J., Fachinetti, D., Bermejo, R., Cocito, A., Costanzo, V., and Lopes, M. (2012). Topoisomerase i poisoning results in PARP-mediated replication fork reversal. Nat. Struct. Mol. Biol. 19(4), 417–423.

Saldivar, J.C., Cortez, D., and Cimprich, K.A. (2017). The essential kinase ATR: Ensuring faithful duplication of a challenging genome. Nat. Rev. Mol. Cell Biol. 18(10), 622–636.

Saldivar, J.C., Hamperl, S., Bocek, M.J., Chung, M., Bass, T.E., Cisneros-Soberanis, F., Samejima, K., Xie, L., Paulson, J.R., Earnshaw, W.C., et al. (2018). An intrinsic S/G2 checkpoint enforced by ATR. Science 361(6404), 806–810.

Sanjiv, K., Hagenkort, A., Calderón-Montaño, J.M., Koolmeister, T., Reaper, P.M., Mortusewicz, O., Jacques, S.A., Kuiper, R. V., Schultz, N., Scobie, M., et al. (2016). Cancer-Specific Synthetic Lethality between ATR and CHK1 Kinase Activities. Cell Rep. 14(2), 298–309.

Schoonen, P.M., Kok, Y.P., Wierenga, E., Bakker, B., Foijer, F., Spierings, D.C.J., and van Vugt, M.A.T.M. (2019). Premature mitotic entry induced by ATR inhibition potentiates olaparib inhibition-mediated genomic instability, inflammatory signaling, and cytotoxicity in BRCA2-deficient cancer cells. Mol. Oncol. 13(11), 2422–2440.

Shiotani, B., and Zou, L. (2009). Single-Stranded DNA Orchestrates an ATM-to-ATR Switch at DNA Breaks. Mol. Cell 33(5), 547–558.

Teloni, F., Michelena, J., Lezaja, A., Kilic, S., Ambrosi, C., Menon, S., Dobrovolna, J., Imhof, R., Janscak, P., Baubec, T., et al. (2019). Efficient Pre-mRNA Cleavage Prevents Replication-Stress-Associated Genome Instability. Mol. Cell 73(4), 670–683. e12.

Toledo, L.I., Murga, M., Zur, R., Soria, R., Rodriguez, A., Martinez, S., Oyarzabal, J., Pastor, J., Bischoff, J.R., and Fernandez-Capetillo, O. (2011). A cell-based screen identifies ATR inhibitors with synthetic lethal properties for cancer-associated mutations. Nat. Struct. Mol. Biol. 18(6), 721–727.

Toledo, L.I., Altmeyer, M., Rask, M.B., Lukas, C., Larsen, D.H., Povlsen, L.K., Bekker-Jensen, S., Mailand, N., Bartek, J., and Lukas, J. (2013). ATR prohibits replication catastrophe by preventing global exhaustion of RPA. Cell 155(5), 1088–1103.

Vendetti, F.P., Lau, A., Schamus, S., Conrads, T.P., O’Connor, M.J., and Bakkenist, C.J. (2015). The orally active and bioavailable ATR kinase inhibitor AZD6738 potentiates the antitumor effects of cisplatin to resolve ATM-deficient non-small cell lung cancer in vivo. Oncotarget 6(42), 44289–44305.

Wallez, Y., Dunlop, C.R., Johnson, T.I., Koh, S.B., Fornari, C., Yates, J.W.T., Bernaldo de Quirós Fernández, S., Lau, A., Richards, F.M., and Jodrell, D.I. (2018). The ATR inhibitor AZD6738 synergizes with gemcitabine in vitro and in vivo to induce pancreatic ductal adenocarcinoma regression. Mol. Cancer Ther. 17(8), 1670–1682.

Yazinski, S.A., Comaills, V., Buisson, R., Genois, M.M., Nguyen, H.D., Ho, C.K., Kwan, T.T., Morris, R., Lauffer, S., Nussenzweig, A., et al. (2017). ATR inhibition disrupts rewired homologous recombination and fork protection pathways in PARP inhibitor-resistant BRCA-deficient cancer cells. Genes Dev. 31(3), 318–332.

You, Z., Kong, L., and Newport, J. (2002). The role of single-stranded DNA and polymerase α in establishing the ATR, Hus1 DNA replication checkpoint. J. Biol. Chem. 277(30), 27088–27093.

Zellweger, Ralph and Lopes, M. (2018). Dynamic Architecture of Eukaryotic DNA Replication Forks In Vivo, Visualized by Electron Microscopy. Methods Mol Biol. 1672, 261–294.

Zellweger, R., Dalcher, D., Mutreja, K., Berti, M., Schmid, J.A., Herrador, R., Vindigni, A., and Lopes, M. (2015). Rad51-mediated replication fork reversal is a global response to genotoxic treatments in human cells. J. Cell Biol. 208(5), 563–579.

Zou, L., and Elledge, S.J. (2003). Sensing DNA damage through ATRIP recognition of RPA-ssDNA complexes. Science 300(5625), 1542–1548.

